# Annual Cultivated Extent and Agricultural Land Use Expansion across the Central Grasslands of North America, 1996-2021

**DOI:** 10.1101/2025.03.18.643874

**Authors:** Sean Carter, Scott L. Morford, Jason Tack, Tyler Lark, Nazli Uludere Aragon, Brady W. Allred, Dirac Twidwell, David E. Naugle

## Abstract

Accurate monitoring of cropland dynamics in North American grasslands is essential for assessing biodiversity threats, guiding sustainable land management, and ensuring food security amid rapid environmental change. We developed a 30-meter resolution dataset capturing annual ‘active’ and ‘cumulative’ cropland (1996 to 2021) across the central grasslands of North America using an Attention U-Net convolutional neural network. Our bias-corrected estimates reveal that while active cropland grew by only 3% (+3.51 ± 1.32 million ha, p > 0.1), the cumulative cropland footprint expanded by 17% (+20.64 ± 0.93 million ha, p < 0.05) relative to the baseline (1996–2000), reaching 142.21 ± 4.84 million ha by 2021. This divergence indicates substantial new conversion or recultivation of previously restored grasslands, occurring at a consistent rate of 0.98 ± 0.04 million ha per year (p < 0.001). Mexico showed the largest relative gain in cumulative cropland area, expanding by nearly half (48%, 1.69 ± 0.06 million ha, p < 0.05). By distinguishing between active and cumulative cropland extents, our dataset enables differentiation between short-term, intermittent cultivation and longer-term land-use legacies, allowing for more nuanced assessments of agriculture’s cumulative effects on biodiversity and ecosystem services at the biome scale. This approach provides critical information for conservation planning and sustainable land management across North American grasslands.

## Introduction

Agricultural expansion, while necessary for global food security, threatens biodiversity and ecosystem services in grassland ecosystems worldwide (Zabel et al, 2019; Scholtz & Twidwell, 2022). In North America’s central grasslands, this expansion has driven large-scale declines in endemic fauna, jeopardizing the ecological integrity of the biome (Green et al., 2005; Brennan and Kuvlesky, 2005; Rosenberg et al., 2019). These land-use changes also disrupt vital ecosystem services, including carbon sequestration, water conservation, and soil health, with consequences to regional and global livestock production (Zhang et al., 2007; Power 2010; Foley et al., 2011). Just as the cumulative impact of past land use shapes long-term soil carbon dynamics (DuPont et al., 2010; Sanderman et al., 2017), the cumulative cropland footprint has far-reaching ecological consequences beyond the area actively farmed in any given year. Once tilled, ecosystems experience fundamental changes that persist long after active farming ceases (Guo & Gifford, 2002; Isbell et al., 2019). Therefore, effective conservation requires a comprehensive tracking of both the active and cumulative cultivation footprint.

Existing cropland datasets provide valuable information but fall short of capturing the full extent of these cumulative impacts. The Global Cropland Expansion dataset (Potapov et al., 2022) offers comprehensive global coverage and high spatial resolution, but lacks the annual temporal resolution needed to track year-to-year changes in cultivation patterns. Similarly, the USDA Cropland Data Layer (CDL) provides detailed crop-specific information at high spatial resolution within the United States, but offers limited temporal coverage prior to 2008 and omits grassland regions of North America outside of the United States. Insufficient annual data limits accurate assessment of short-term cultivation dynamics and their legacy impacts.

Consequently, researchers struggle to disentangle the immediate impacts of active cultivation from the broader, long-term consequences of the cumulative cropland footprint, impeding efforts to evaluate the sustainability of agricultural expansion, especially onto lands that are marginal for agricultural production (Johnston, 2014; Wright & Wimberly, 2013; Lark et al., 2015; Mladenoff et al., 2016; Lark et al., 2020).

The ecological impact of agricultural expansion is evident in the widespread declines in grassland birds, mammals, and insects (Herkert, 1994; Greer et al., 2016; Ceballos et al., 2010; Rosenberg et al., 2019; Sánchez-Bayo & Wyckhuys, 2019), signaling a biome under intense pressure. While the active cropland extent contributes to these declines, the legacy effects of past cultivation may be equally important, yet harder to quantify. Understanding these cumulative impacts is crucial for predicting the ecological trajectories of abandoned or idled agricultural lands and for assessing the vulnerability of remaining grasslands to future conversion. High-resolution data that capture both active and cumulative cropland extents are therefore essential for effective conservation planning (Tack et al., 2023).

Recent advancements in deep learning and computer vision offer powerful tools for extracting complex spatial patterns from satellite imagery (Zhang et al., 2016; Ma et al., 2019; Kussul et al., 2017). Here, we introduce the Annual Cultivated Extent (ACE) dataset, a 30-m annual dataset capturing cropland extent across North American central grasslands from 1996 to 2021. By integrating Landsat imagery (USGS, 2023) with vegetation data from the Rangeland Analysis Platform (Allred et al., 2021) and applying deep learning, ACE provides annual estimates of both “active” and “cumulative” cropland areas. In this context, “active croplands” are areas under cultivation during a given year, while “cumulative croplands” include both the actively cultivated areas for that year, and any land cultivated in any previous year of the study period. ACE helps fill a critical knowledge gap about the full ecological impact of agricultural land use change across the largest contiguous grassland biome in North America.

## Methods

### Overview

This study generated annual 30-m resolution maps of active and cumulative cropland extent across central North America from 1996 to 2021. Cropland classification was performed using an Attention U-Net convolutional neural network (Oktay et al., 2018), trained on Landsat spectral indices and Rangeland Analysis Platform vegetation data (USGS 2023; Allred et al., 2021). Our training dataset combined manually labeled image chips with synthetic labels derived from existing products (Potapov et al., 2022). To enhance data quality, we implemented post-processing techniques to mitigate cloud cover and data gaps. Finally, bias correction using a stratified reference sample (Stehman 2014; Oloffson et al., 2014) was applied to refine area estimates. Our complete workflow is depicted in Figure 1.

**Figure 1:**
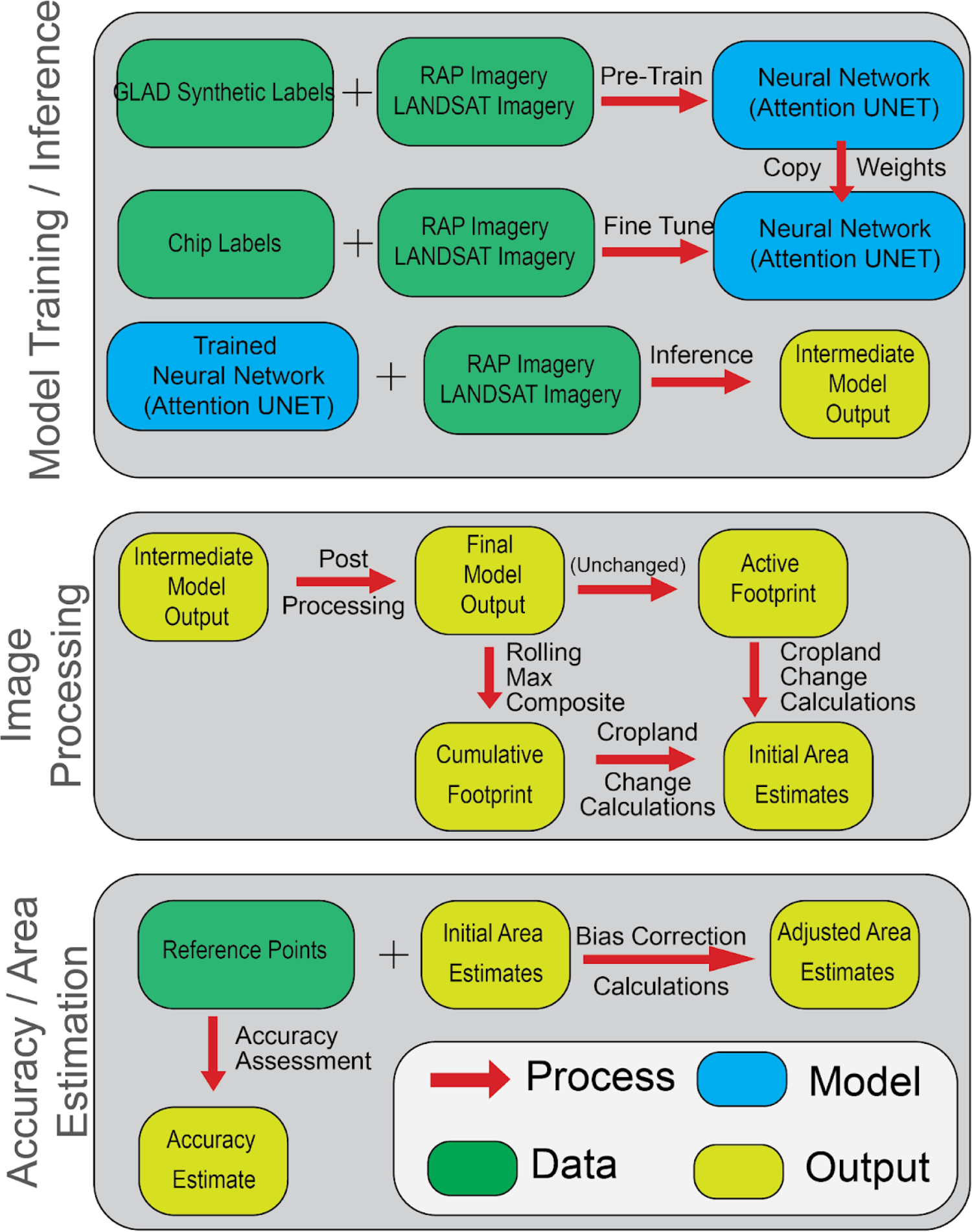
Workflow diagram illustrating data sources, processing steps, and analysis pipeline used to create the Annual Cropland Extent (ACE) dataset.

### Study Area

Our study area focuses on the central grasslands of North America (Figure 2), a biome of critical importance for biodiversity and ecosystem services, including vital habitat for migratory bird populations experiencing declines linked to habitat loss (Brennan & Kuvlesky 2005, Pool et al., 2014; Rosenberg et al., 2019). Specifically, we examined portions of the Great Plains, North American Deserts, and Southern Semi-Arid Highlands ecoregions within the North American Central Flyway (La Sorte et al., 2014, Omernik and Griffith, 2014). This region’s importance for agriculture and biodiversity conservation makes it an ideal location to study the interplay between agricultural land use and ecological change across a broad range of biophysical, socioeconomic, and geopolitical contexts.

**Figure 2:**
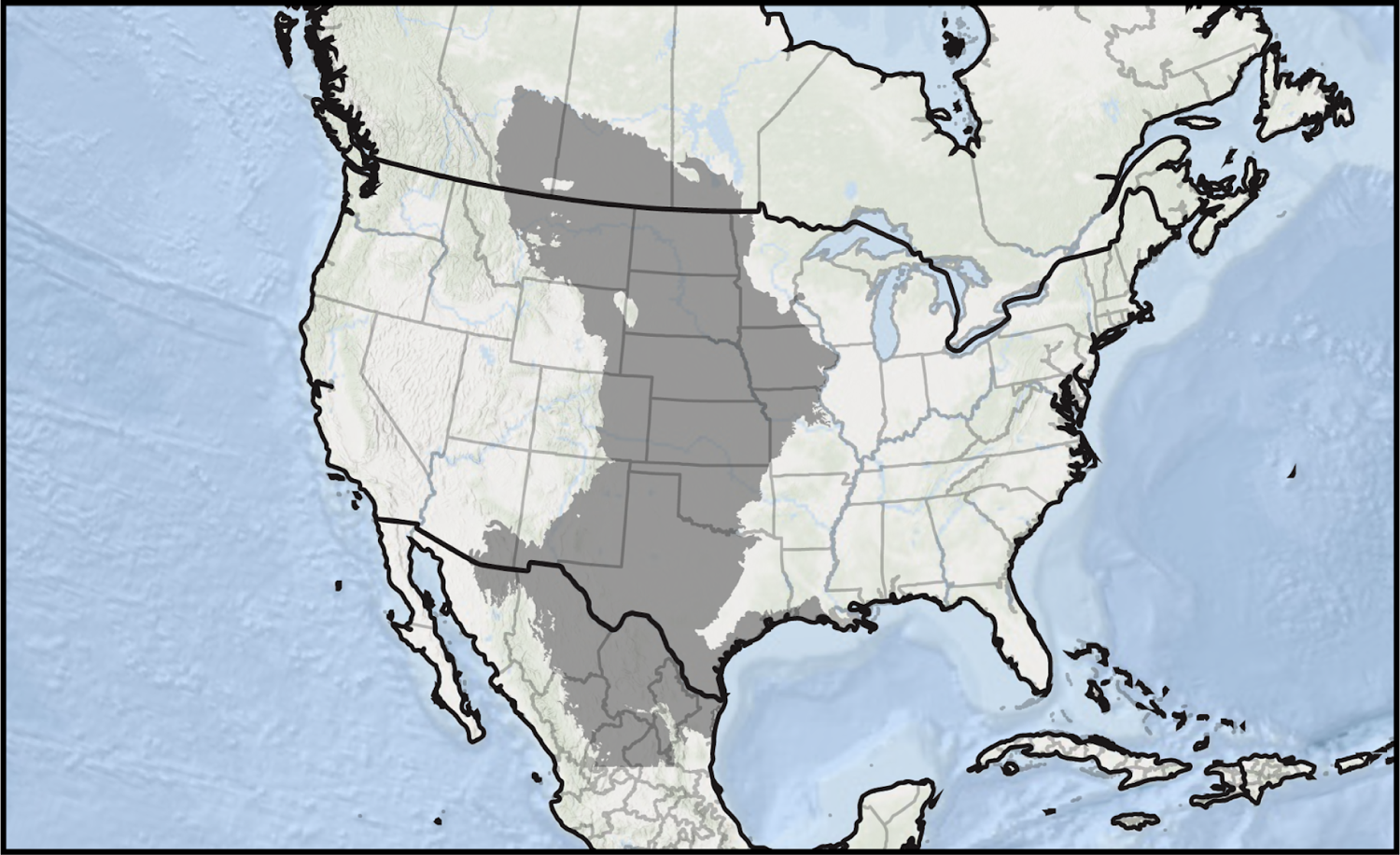
Cropland modeling in this analysis (shown in gray) encompassed the central grasslands of North America, spanning Canada, Mexico, and the United States, and overlapping with central grassland bird conservation regions.

### Data Sources

#### Input Features

We derived spectral indices from Landsat Collection 2 surface reflectance Tier 1 data (Crawford et al., 2023) and combined them with annual fractional vegetation cover data from the Rangeland Analysis Platform (Allred et al., 2021; RAP). Using Google Earth Engine (GEE), we created cloud-free, seasonally composited Landsat Mosaics for each year. Cloud cover was masked using the Landsat Quality Assessment (QA) bands. To maximize the contrast between cropland and non-cultivated vegetation, we focused on the senescence period of natural vegetation (day-of-year 244-305), when actively growing crops are more readily distinguishable. We calculated three spectral indices: Normalized Difference Vegetation Index (NDVI), Normalized Burn Ratio (NBR), and Normalized Difference Moisture Index (NDMI), chosen for their sensitivity to vegetation characteristics and known utility in cropland mapping (Vidican et al., 2023). From RAP, we extracted annual fractional cover layers for annual forbs and grasses (AFG), perennial forbs and grasses (PFG), and tree cover (TRE). RAP fractional cover layers were extended from the U.S footprint to Canada and northern Mexico to include the entirety of our study area. These layers provide valuable information on vegetation composition and structure, aiding in the differentiation of croplands from other land cover types. The final input features for our model consisted of pixel-wise maximum composites for each Landsat index, stacked with corresponding annual RAP fractional cover layers.

#### Training Targets

Model targets were derived from two labeling sources to ensure comprehensive representation of cropland characteristics across the study area. First, we manually annotated 30 disparate 640,000 hectare scenes within 6 different years (in total, 19.2 million hectares) in areas of the southern Great Plains and the Mexican desert regions where existing cropland products consistently underperformed. This targeted approach allowed us to improve model accuracy in challenging areas with heterogeneous landscapes and fragmented agricultural fields. An analyst visually interpreted agricultural fields using true-color and false-color composites from Landsat and Google Earth Imagery (hereafter, the “chip” dataset). Second, for the northern extent of our study area, where existing cropland identification has historically been more reliable, we leveraged the Global Land Analysis and Discovery (GLAD) cropland extent layer (Potapov et al., 2022). We randomly sampled GLAD data 30,000 times (years 2003, 2007, 2011, 2015 and 2019) and combined it with 5000 samples from the chip dataset (years 1995, 1997, 1999, 2003, 2005, and 2011) to create a balanced and geographically diverse training dataset across 9 years. Each sample consisted of a potentially overlapping 512 x 512-pixel tile (∼2,400 hectares), with labels sourced either from our manual chip annotations or from the synthetic GLAD dataset. The chip dataset was augmented through four 90-degree rotations to increase training data diversity and improve model generalization. This process yielded approximately 200 GB of GLAD data and 30 GB of chip data. We reserved 20% of the combined dataset for validation during model training.

### Model Training and Inference

We employed the Attention U-Net architecture (Ronneberger et al., 2015; Oktay et al., 2018) for the semantic segmentation of cropland (i.e., classifying each pixel as cropland or non-cropland). Such segmentation models have been shown to better characterize spatial heterogeneity of target classes and differentiate based on structural attributes relative to pixel-based approaches (Kussul et al., 2017). To prevent overfitting and improve generalization on unseen data, we implemented a four-stage training strategy (Supplemental 3.1).

Following model training, we generated annual cropland predictions for 1995 - 2021. For each year, we applied the same cloud filtering and seasonal compositing methodology used for the training data. These annual composites were exported from GEE and processed in tiled segments to facilitate efficient inference using the trained Attention U-Net model. The resulting intermediate predictions were then seamlessly stitched together and uploaded back to GEE as an image collection.

### Image Processing

We then applied two key post-processing steps:

1. *Woodland Masking:* To address commission errors, particularly the misclassification of woodlands as cropland, we applied a forest mask developed using a Random Forest model trained on NDVI, Gross Primary Productivity, precipitation data (Hengl & Parente, 2022), RAP data, and NLCD labels (see Supplemental 3.2 for details). This model, trained with 550 points and achieving 93% accuracy on a validation set, specifically improved forest detection within where NLCD often underperforms (Wang & Mountrakis, 2023). The resulting mask reclassified pixels identified as both woodland and cropland to woodland across the conterminous United States.
2. *Gap filling*: We addressed data gaps caused by cloud cover or image acquisition issues using a two-year moving window temporal filter. For each year *t* from 1996–2021, we generated a pixel-wise maximum composite from the predictions for years *t* and *t*-1. This approach filled small gaps under the assumption of persistent cropland status between consecutive years. This post-processing produced a time series of 26 annual cropland maps (1996–2021).

### Cropland change calculations

We quantified changes in both active cropland (areas classified as cropland in a specific year) and cumulative cropland (area cropped at any time up to a given year). Active cropland was presented by a binary map *P*_*t*_(*x*, *y*), for each year *t,* where:

*P*_*t*_ (*x*, *y*) = 1 if the pixel is cropland

*P*_*t*_ (*x*, *y*) = 0 otherwise

Cumulative cropland,*C*_*t*_ (*x*, *y*), up to year *t*, was calculated as:

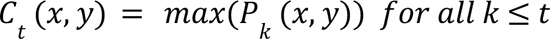

Annual cropland area (*A*_*t*_) was calculated by summing the area of all cropland pixels:

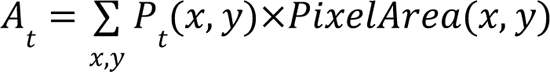

### Accuracy and Area Estimation

We assessed model accuracy and adjusted area estimates for bias using a stratified random sampling approach. We generated 270 reference samples annually from 1996 to 2021. Stratified sampling ensures adequate representation of different land-use and land-cover types, leading to a more robust accuracy assessment (Olofsson et al., 2014). Based on observed discrepancies in model performance likely related to differences in training data availability and landscape characteristics, we divided the study area into two subregions: the Great Plains and the Southern Subregion (comprising the Southern Semi-Arid Highlands, North American Deserts, and Temperate Sierras ecoregions; Omernik and Griffith, 2014).

Following protocol of Potapov et al. (2022), we generated a reference sample across five strata designed to capture potential temporal, spatial, and pattern-based errors (see Supplemental 3.3 for detailed strata definitions). These strata included stable cropland, cropland gain, cropland loss, areas with uncertain predictions, and stable non-cropland. To account for regional differences in area estimation within the Southern Subregion, where cropland patches are smaller and more dispersed compared to the Great Plains, we employed an adaptive buffer to contiguous cropland sections in the Southern Subregion. Specifically, we analyzed the spatial autocorrelation of cropland fields in the Great Plains using a variogram to determine a characteristic field-neighborhood distance, which was then applied as a buffer around cropland patches in the Southern Subregion. For example, a 40-hectare cropland field in the Southern Subregion was assigned an 18 km buffer zone, which then defined the reference sampling frame for that field to better capture the local agricultural landscape. This process was repeated for each contiguous cropland field in the Southern Subregion. This approach was necessary to ensure reliable area estimates in the Southern Subregion, where the large proportion of non-agricultural land would otherwise lead to unstable bias corrections when calculating cropland area. Each year’s reference sample included 200 points from the Great Plains and 70 from the Southern Subregion, reflecting the relative sizes of the two subregions.

A visual analyst classified each reference sample as “cropland” or “non-cropland” using RAP and Landsat imagery for the target year.

### Accuracy Metrics and Area Calculation

We calculated standard accuracy metrics— overall accuracy (*OA*), and *F*1 score—for each subregion, where each stratum (*h*) is weighted by its proportional area (*w*_*h*_):

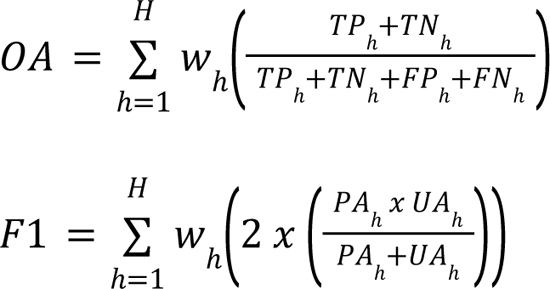

Where:

*H* is the number of strata

*w*_*h*_ is the proportion of the total area occupied by stratum *h*

*TP*_*h*_ represents true positives (correctly classified cropland)

*TN*_*h*_ represents true negatives (correctly classified non-cropland)

*FP*_*h*_ represents false positives (non-cropland incorrectly classified as cropland)

*FN*_*h*_ represents false negatives (cropland incorrectly classified as non-cropland)

*PA*_*h*_ represents producer’s accuracy;

*TP*_*h*_ / (*TP*_*h*_ +*FN*_*h*_)

*UA*_*h*_ represents user’s accuracy;

*TP*_*h*_ / (*TP*_*h*_ +*FP*_*h*_)

(All *h* subscripts represent values observed in stratum *h*)

Error-adjusted area estimates for cropland and non-cropland were then computed following Olofsson et al. (2014), accounting for commission and omission errors and weighing by the proportional area of each stratum. The estimated area for each class was calculated as:

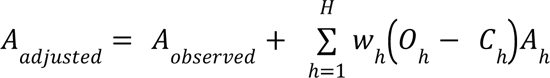

Where:

*A*_*adjusted*_ is the bias-adjusted cropland area

*A*_*observed*_ is the originally observed cropland area

*H* is the number of strata

*w*_*h*_ is the proportion of the total area occupied by stratum *h*

*O*_*h*_ is the omission error rate for stratum *h*

*C*_*h*_ is the commission error rate for stratum *h*

*A*_*h*_ is the area of stratum *h*

The standard error for *A*_*adjusted*_ was calculated using:

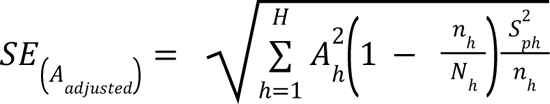

Where:

*A*_*h*_ is the originally observed cropland area

*n*_*h*_ is the sample size in stratum *h*

*N*_*h*_ is total number of possible sampling units in stratum *h*_2_

*S*^2^_*ph*_ is the sample variance for stratum *h*, calculated as *S*^2^_*ph*_ = *p*_*−h*_ (1 − *p*_−*h*_)

*p* is the mean cropland change proportion of samples in stratum *h*

We applied this methodology independently to each subregion and then combined the results to generate final area estimates and associated uncertainties for the entire study area. We applied the above methodology annually, resulting in adjusted area and error estimates for both cumulative and active footprint statistics.

To account for intermittent croplands returning to cultivation in early years of our time series, we defined a baseline period (1996-2000) that establishes an initial extent of cumulative crop footprint. This approach helps reduce the influence of historic intermittent cropping or cyclical longer-term fallowing periods that would otherwise upwardly bias our estimates of cumulative cropland extent in early years of the study period. For example, our product does not capture conditions prior to 1996, so change estimates relative to the earliest time point would be biased towards an overrepresentation of recurrent agricultural lands that undergo cyclic fallow periods and have been in production prior to our time window. Subsequent change estimates were calculated relative to this baseline.

To understand cropland change, we applied an ordinary least squares regression to values observed between 2000 and 2021, thus accounting for the baseline observations. We calculated the overall change statistics by multiplying the slope by the time period, with the associated uncertainty derived from the standard error of the slope estimate. This approach allowed us to characterize both the magnitude and statistical significance of agricultural land use changes while accounting for year-to-year variability in the estimates.

## Results

We produced annual layers of active and cumulative cropland extent for the central grasslands of North America from 1996 to 2021, covering an area of 349.6 million ha across Canada, Mexico, and the United States. The ACE dataset is available for download from the Zenodo Open Research Repository (https://zenodo.org/records/14607083) and as an interactive map application in Google Earth Engine (*see Data Availability*). The ACE dataset includes 26 annual GeoTIFF rasters, each containing two bands that represent active and cumulative cropland, respectively.

The cropland maps achieved an Overall Accuracy and F1 score of 0.83 and 0.89, respectively, when compared to our independently collected reference dataset. The Great Plains subregion had an Overall Accuracy of 0.82 and an F1 score of 0.88, and the Southern Subregion reported an Overall Accuracy of 0.86 and an F1 score of 0.90. Examples of cropland classification from our model are provided in Figure 3.

**Figure 3:**
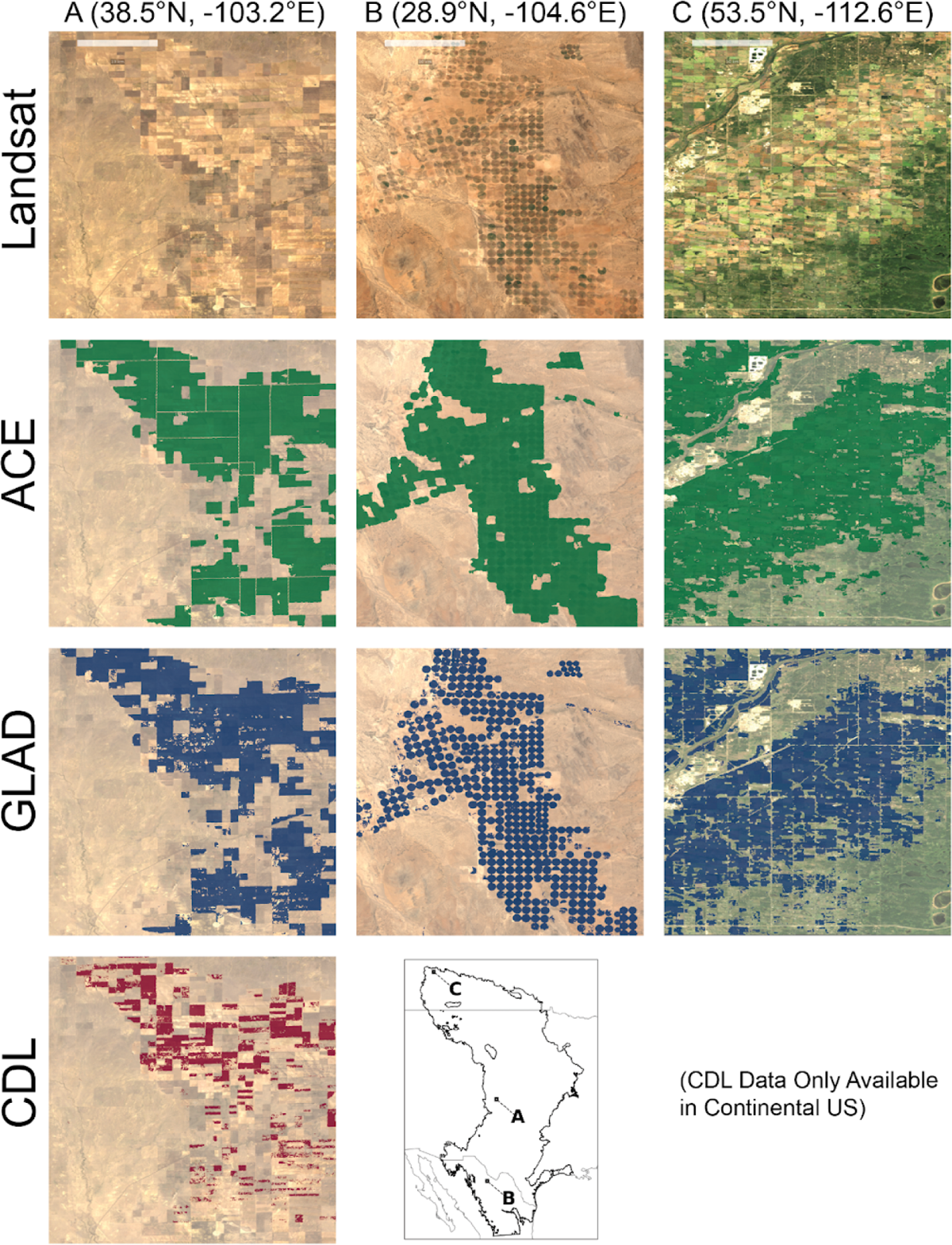
Model comparison with existing data products. Landsat imagery is an annual median composite of Landsat 5 and 7 Level 2 Surface Reflectance. ACE: Annual Cropland Extent Dataset from this paper; GLAD: Global Cropland Extent product (Potapov et al., 2022). CDL: Cropland Data Layer (USDA 2023). CDL data only available in continental United States

Using our bias-correction methods, we identified a cumulative cropland footprint of 142.21 ± 4.84 million ha across the study region (Table 1). Compared to the baseline period (1996 - 2000), we observed that between 2000 and 2021, the cumulative cropland area increased by 20.64 ± 0.93 million hectares (16.99%, p < 0.05). Over this same period, there was little evidence that the active cropland area (land under cultivation in a given year) increased (3.51 ± 1.32 million ha; 3.32%, p > 0.1), remaining relatively stable over the entire study period (1996 - 2021, Figure 4). The cumulative cropland area was observed to increase linearly at a rate of 0.98 ± 0.04 million hectares per year (p < 0.001) between 2000 and 2021.

**Figure 4:**
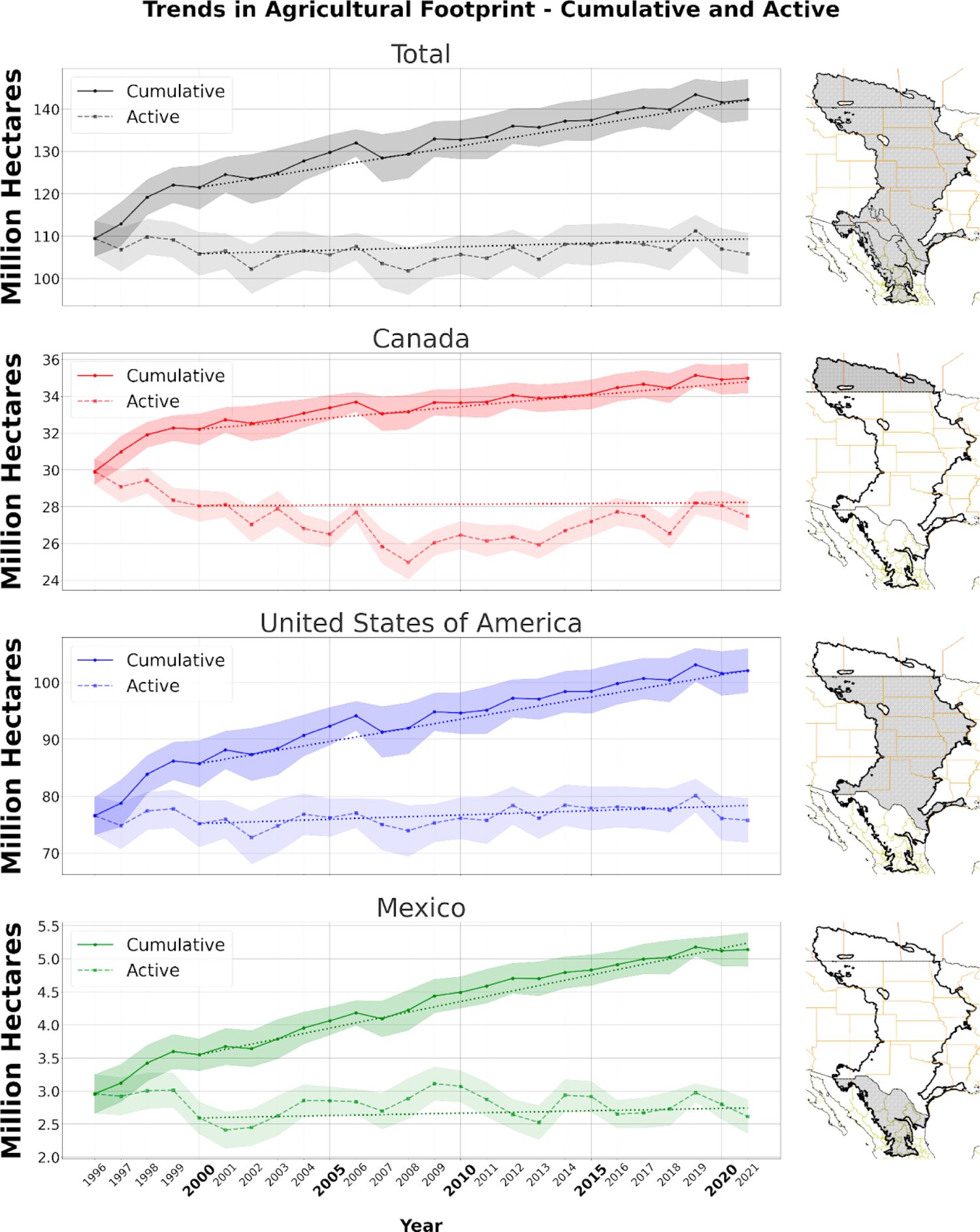
Bias adjusted cumulative and active cropland change estimates in Canada, the United States, and Mexico from 1996 to 2021. Solid lines represent the cumulative cropland footprint. Dashed lines show the active cropland area. Error bars indicate the 95% confidence interval for area estimates. Dotted line represents least squares regression.

**Table 1.** Bias-adjusted active and cumulative cropland area and changes (2000–2021) across the study area (in millions of hectares)

| Region(s) | Cropland Metric | ROI Area | Cropland: Year 2000 | Cropland: Year 2021 | Cropland Change | Percent Change |
| --- | --- | --- | --- | --- | --- | --- |
| Canada | Active | 46.0 | 28.0 ± 0.8 | 27.5 ± 0.8 | 0.2 ± 0.7† | 0.65% |
| Canada | Cumulative |  | 32.2 ± 0.8 | 35.00 ± 0.8 | 2.58 ± 0.16* | 8.02% |
| Mexico | Active | 67.7 | 2.59 ± 0.24 | 2.61 ± 0.25 | 0.15 ± 0.13 | 5.80% |
| Mexico | Cumulative |  | 3.55 ± 0.24 | 5.14 ± 0.25 | 1.69 ± 0.06* | 47.60% |
| United States | Active | 244.8 | 75.2 ± 4.1 | 75.8 ± 3.7 | 3.2 ± 1.0† | 4.23% |
| United States | Cumulative |  | 85.7 ± 4.1 | 102.1 ± 3.9 | 16.37 ± 0.75* | 19.10% |
| **All** | **Active** | **358.4** | **105.8 ± 5.1** | **105.9 ± 4.8** | **3.5 ± 1.3** | **3.32%** |
| **All** | **Cumulative** |  | **121.5 ± 5.1** | **142.2 ± 4.8** | **20.64 ± 0.93\*** | **16.99%** |

\*: $p < 0.05$ , †: $0.05 \leq p < 0.1$ ; Mann-Kendall Test; ROI: Region of Interest; Baseline cumulative cropland area for the year 2000 represents bias-corrected cumulative cropland area calculated in 2000. Values reported in millions of hectares; error estimates represent 95% confidence intervals.

Statistically significant increases in cumulative cropland area were observed in all three countries, while active cropland patterns were more temporally variable and showed no pronounced trends (Table 1, Figure 4). Between 2000 and 2021, cumulative cropland expanded at annual rates of 0.12 ± 0.01 million ha for Canada, 0.08 ± 0.003 million ha for Mexico, and 0.78 ± 0.05 million ha in the United States. In Canada, active croplands declined from 1996 to 2008 and then increased, resulting in little net change between 2000 and 2021, while those in the United States showed modest increases from the early 2000s and late 2010s.

A comparison of our active and cumulative cropland area estimates with existing national and spatial datasets is presented in Table 2. For Canada, our active cropland estimates between 1996 and 2021 closely matched Statistics Canada field inventories, with a maximum difference of 2.9%. However, North American Land Change Monitoring System (NALCMS) cumulative cropland estimates were 22% lower than ours and even lower than Statistics Canada’s 2020 active cropland area. In Mexico, our cumulative cropland estimates and those from INEGI were largely consistent, differing by at most 10.3% (Instituto Nacional de Estadística y Geografía, 2020). For the United States, active cropland area estimates between ACE and the Cropland Data Layer (CDL) differed by up to 17.0% between 2012 and 2021. Our model indicated a slight decline in active cropland during this period, while the CDL showed an increase. Both datasets, however, revealed similar directional trends for cumulative cropland area, although the CDL reported a larger magnitude of change.

**Table 2:** Comparison of modeled cropland estimates to national statistics (in millions of hectares)

| Country | National Estimate | Cropland Metric | Time Period | ROI Adjustment | Comparison |  |
| --- | --- | --- | --- | --- | --- | --- |
|  |  |  |  |  | National Product | This Analysis |
| Canada | StatCan<sup>[a]</sup> | Active | 1996-2021 | 0.88 multiplier<sup>1</sup> | 30.0 → 28.3 | 29.9 ± 0.7 → 27.5 ± 0.8 |
| Canada | NALCMS<sup>[b]</sup> | Cumulative | 2020 only | ROI overlap<sup>2</sup> | 28.0 | 35.0 ± 0.8 |
| Mexico | INEGI<sup>[c]</sup> | Active | 2000-2020 | - | - | 2.5 ± 0.2 → 2.8 ± 0.2 |
| Mexico | INEGI<sup>[c]</sup> | Cumulative | 2000-2020 | Direct comparison<sup>4</sup> | 3.4 → 4.6 | 3.6 ± 0.2 → 5.1 ± 0.2 |
| United States | CDL<sup>[d]</sup> | Active | 2012-2021 | ROI overlap<sup>3</sup> | 66.1 → 73.2 | 78.4 ± 3.3 → 75.8 ± 3.9 |
| United States | CDL<sup>[d]</sup> | Cumulative | 2012-2021 | ROI overlap<sup>3</sup> | 86.85 → 96.79 | 97.2 ± 3.3 → 102.1 ± 3.9 |

Notes: <sup>1</sup>StatCan multiplier based on average percent of croplands in the Region of Interest (ROI); <sup>2</sup>Single-year comparison (NALCMS; 2020 compared with 2021 model) aligned with study area overlap; <sup>3</sup>Comparison restricted to overlapping areas within the ROI in the same years; <sup>4</sup>INEGI area does not overlap completely with CropScope ROI so direct values were compared. Data sources: <sup>a</sup>Statistics Canada (2023), <sup>b</sup>Latifovic et al. (2020), <sup>c</sup>INEGI (Instituto Nacional de Estadística y Geografía, 2020); <sup>d</sup>USDA-NASS (2023). ROI: region of interest; StatCan: Statistics Canada; NALCMS: North American Land Change Monitoring System; INEGI: Instituto Nacional de Estadística y Geografía; CDL: Cropland Data Layer; '-' indicates no data.

## Discussion

The ACE dataset offers a unified resource to evaluate cropland changes across North America’s central grasslands from 1996 to 2021. Unlike epoch-based products that lack annual resolution (e.g., GLAD) or those limited by political boundaries to a single country (e.g., CDL), ACE provides consistent annual coverage across the entire biome regardless of national borders. This allows for analysis of long-term trends and cross-border dynamics that was previously not possible with existing datasets that lacked the necessary spatial or temporal coverage. Our ACE dataset is also among the first to quantify agricultural change in Mexican grasslands and deserts, a region often overlooked in analyses of North American grasslands. Lastly, by predicting both active and cumulative cropland extent, ACE allows users to now differentiate the effects of short-term, intermittent cultivation from longer-term land-use trends and legacies. This distinction is crucial for modeling ecological impacts, allowing for more nuanced assessments of agriculture’s cumulative effects on biodiversity, soil health, and carbon storage at the biome scale.

ACE helps reveal dynamic changes in the distribution of active croplands over the past two-plus decades (Figure 5, left panel). In the northern Great Plains, active cropland areas have increased in the United States but remained stable or declined in Canada, potentially reflecting differences in agricultural policies, economic drivers, or shifting weather patterns (Hatfield & Beres, 2019; Gerber et al., 2024). In the central and southern Great Plains, patterns are more variable, likely influenced by local factors such as water limitations from the High Plains aquifer or the expansion of urban areas (Xie et al., 2024; Scanlon et al., 2012; Rhodes et al., 2023). The high temporal resolution of ACE also captures shorter-term regional shifts (Figure 5, right panels). For example, in northern Missouri (United States), active cropland expanded from 1996 to 2000 and then declined from 2011 to 2015, mirroring state-level trends (USDA 2023).

**Figure 5:**
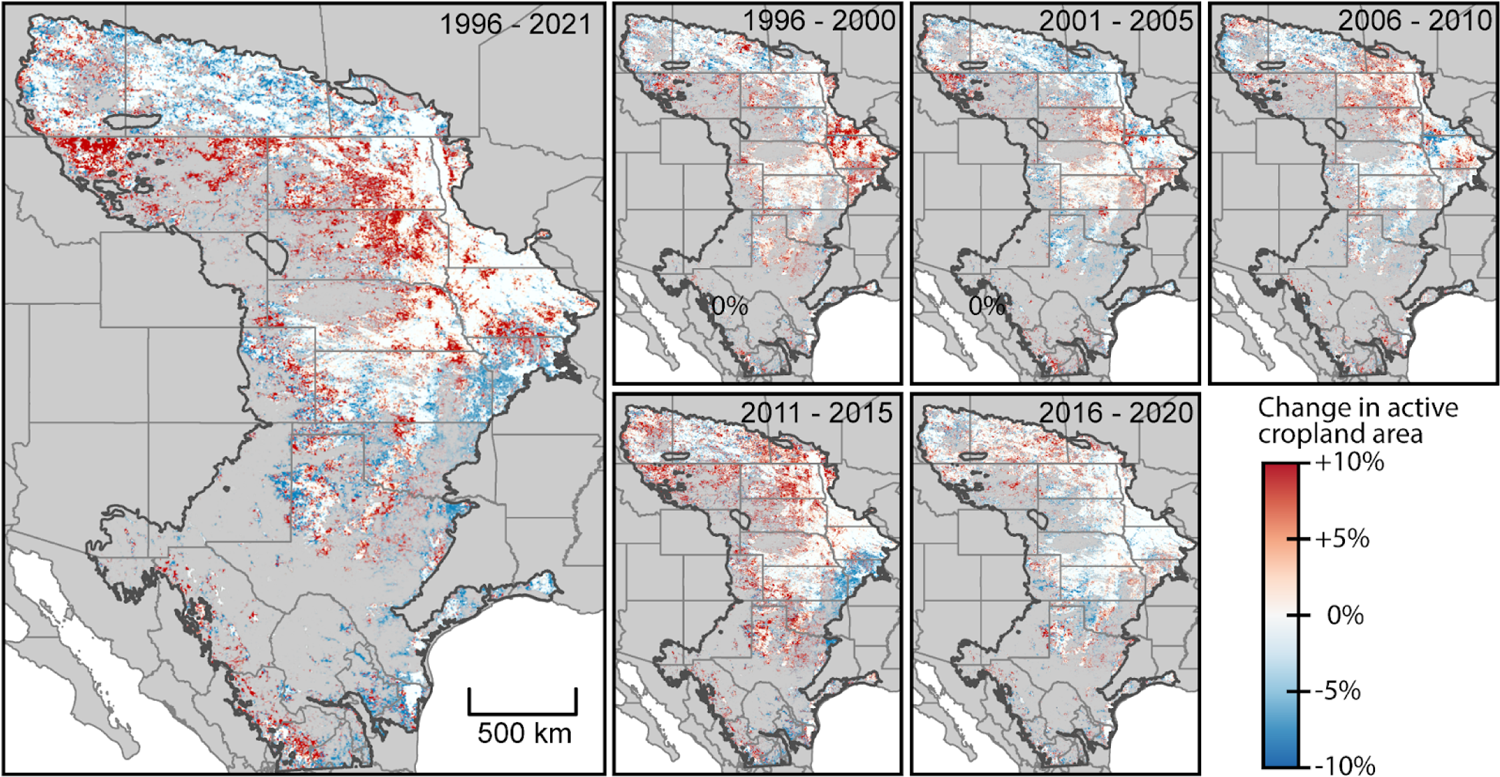
Changes in active cropland area across the study region over the entire study period (left) and five-year epochs (right). Red areas indicate an increase in active cropland during the specified period, blue areas indicate a decrease, and white areas indicate no change. Each 5-km pixel’s opacity reflects the proportion of cropland in that pixel, with more opaque pixels indicating a greater cropland fraction. To minimize overestimation of change due to short-term crop rotations, we compared three-year maximum composites of active cropland centered on the beginning and end years of each five-year epoch. For the 1996-2021 comparison, we used five-year maximum composites from the beginning and end of the time series.

Between 2016 and 2020, active croplands continued to expand in the western marginal lands of the northern Great Plains, while contracting in often-irrigated locations overlying the High Plains Aquifer. These patterns are consistent with recent analyses of North American land-use change (Lark et al., 2020; Xie and Lark, 2021; Xie et al., 2024).

Although Mexico had the smallest cropland footprint in our study area, ACE shows Mexico experienced the greatest relative increase in cumulative cropland area, expanding by nearly 48% from the baseline period (1996–2000) to 2021 (Table 1). Much of this growth occurred in the Mexican states of Chihuahua and northern Durango, aligning with the expansion of large irrigation projects in the Chihuahuan Desert (Ceballos et al., 2005; Pool et al., 2014; Bonilla-Moheno & Aide, 2020; SAGARPA, 2017). We estimate that croplands in Chihuahua expanded at 39,072 hectares per year between 2012 and 2021 (∼400,000 total; Figure 6), consistent with estimates from Pool et al. (2014). In contrast, declines in active cropland were concentrated in the eastern states of Nuevo León and Tamaulipas, where many previously cultivated areas have either transitioned to shrub cover or converted to pastureland.

**Figure 6:**
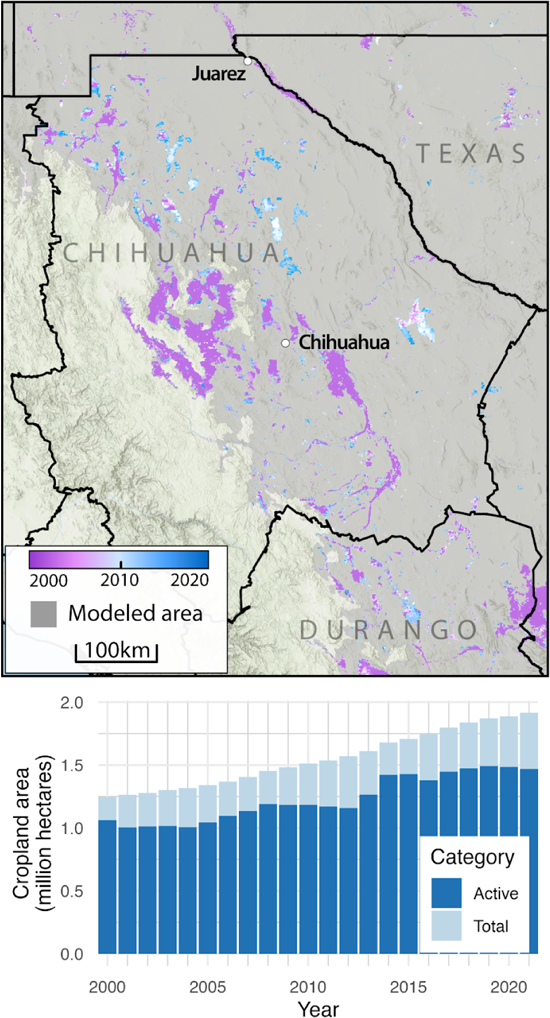
Change in cropland area in Chihuahua, Mexico. Color represents the year cropping was first detected. The model area includes central and eastern Chihuahua, but excludes the western region in the Sierra Madre Occidental, so cropland in valleys of that region will not be represented here.

An inherent limitation of this study lies in its reliance on optical remote sensing, including potential data gaps and classification errors from cloud cover addressed through temporal compositing. In our cropland area calculations, we address such errors using bias-correction methods; however, even these estimates are subject to reference data reliability, and ultimately users should interpret pixel-level results with caution, particularly in regions like the Great Plains where distinguishing cropland from pasture is challenging. To facilitate local-scale exploration, we provide an application for visualizing the annual and time series cropland data (see Data Availability). Importantly, cropland area estimates from ACE align with existing national monitoring programs and data products (Table 2, Figure 3), including independent estimates of cropland expansion in the Great Plains (World Wildlife Fund, 2023). Extending beyond the temporal and spatial coverage of these existing resources, ACE offers unique capabilities for cross-border analyses of land-use change in central North American grasslands.

The ACE dataset reveals a key dynamic. While the cumulative agricultural footprint expands, the area under active cultivation has remained relatively stable year to year during our study period (Table 1, Figure 4). This phenomenon can be attributed to several factors. First, our approach tracks only increases in cumulative cropland footprint, such that permanent removals of cropped areas (e.g. when cropland gets converted to urban or other uses, including abandonment) do not affect its accounting. Thus, for example, active croplands lost to urbanization that are “replaced” or compensated by new croplands elsewhere add to the cumulative footprint but not to the active footprint (Emili and Greene, 2014). Second, there often exists an ongoing churning of agricultural lands, with lower-productivity, so-called “marginal” lands rotating in and out of cultivation over varying short-(e.g. years) to long-term (e.g. decadal) time scales. This reality, paired with the finite duration of our study period, results in our cumulative cropland estimates capturing and characterizing the ever-expanding footprint of cultivated agriculture regardless of whether active cropped area increases, decreases, or stays flat in any given year.

One example conservation implication of our findings is that strategically incentivizing the retirement or restoration of marginal croplands can reduce environmental impacts without significantly affecting agricultural production. These marginal lands often represent some of the most environmentally sensitive landscapes remaining in the Central Grasslands (Lark et al., 2020). Datasets like ACE are crucial to realize this potential by allowing spatial targeting of conservation policy alternatives that are congruent with the biophysical and socioeconomic drivers of marginal land cultivation. However, it’s important to acknowledge that spatial targeting alone faces implementation challenges when broader economic forces favor conversion, regardless of land productivity (Miao et al., 2016). For targeting to succeed at meaningful scales, complementary efforts are needed to enhance the economic competitiveness of grasslands in ways that recognize their inherent values to society and agriculture (Ribaudo et al., 2010).

From a grassland perspective, ACE is particularly valuable for identifying areas with low (temporal) frequency of cultivation (Figure 7a), revealing opportunities for proactive conservation to maintain and grow core grassland areas. By leveraging the agricultural “churn” dynamics described earlier, conservation efforts can target these transitional zones where economic barriers to maintaining grassland cover may be lower. Current prioritization approaches often miss critical areas exhibiting ‘pioneering’ cultivation patterns - isolated grasslands undergoing conversion to cropland, even as other nearby areas show decreasing cultivation frequency (Fig. 7b). Using ACE will enable future prioritizations to incorporate non-stationarity into more reliable predictors of future cropland conversions (e.g., Bedrosian et al., 2024; Olimb & Robinson, 2019; Barnes et al., 2025).

**Figure 7.**
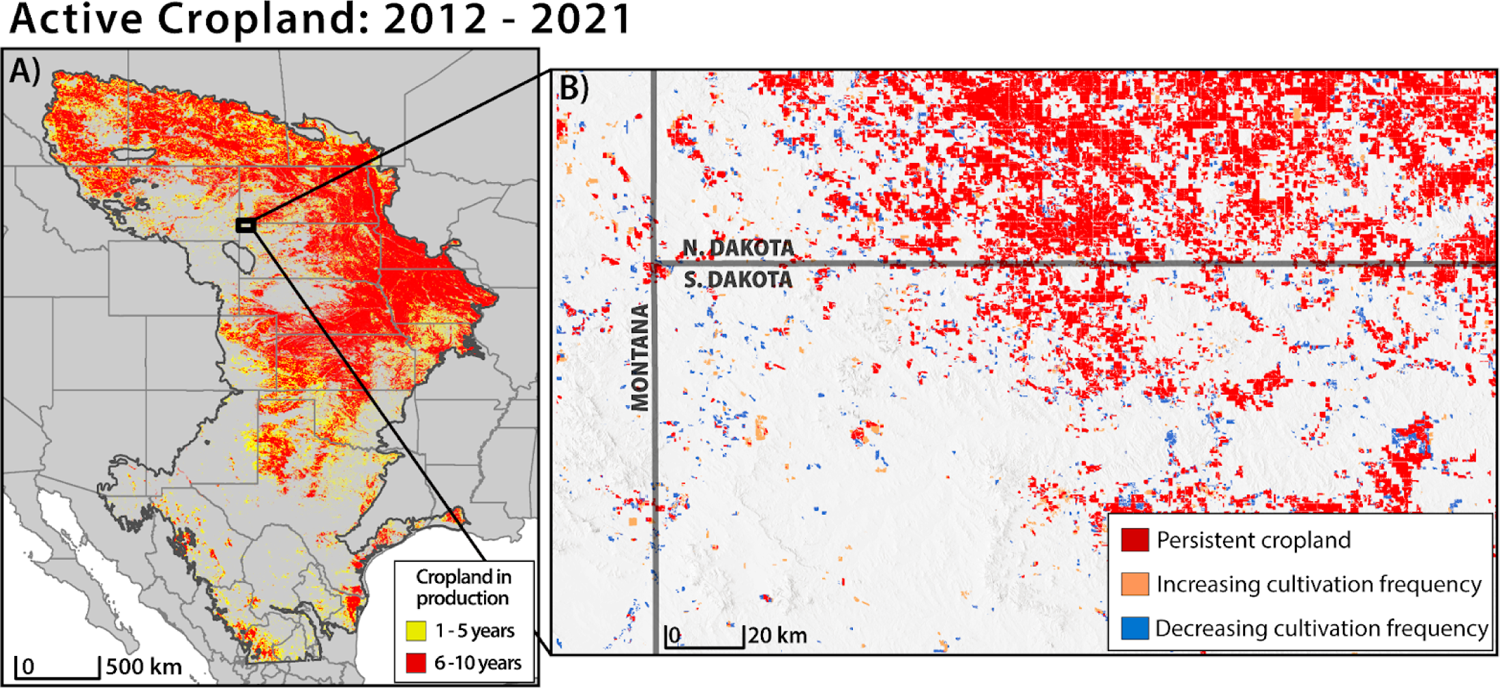
Active cropland dynamics, 2012-2021. Left panel (A): Red areas show persistent cropland (>5 years cultivation), yellow shows intermittent cultivation (≤5 years). Right panel (B): A detailed view of cropland dynamics in the Northern Great Plains illustrates agricultural “churn” in the western most part of South Dakota - red indicates persistent cropland, orange shows cropland with intermittent increasing cultivation frequency, and blue represents intermittent decreasing cultivation frequency. Even as cultivation decreases in some parts of the landscape, areas of increasing cultivation, which includes new grassland conversion, continues nearby, highlighting the importance of spatial and temporal context when prioritizing conservation of intact grassland systems.

Drawing inspiration from collaborative conservation efforts across North American grasslands, ACE data could effectively guide conservation investments by identifying remaining intact areas. The “defend the core” strategy (Doherty et al., 2024) provides a complementary framework that could be applied to strategically invest conservation resources in core grasslands to prevent further degradation. This approach aligns with existing grassroots initiatives like the Great Plains Grassland Initiative, which integrates vegetation data with landowner input to identify priorities. Multi-scale efforts, including the Central Grasslands Roadmap Assessment and Tri-National Grasslands Initiative, have already begun mapping core functional grasslands using threat-based frameworks. ACE data enhances these efforts by providing precise temporal information on cultivation patterns, enabling targeted conservation across the landscape continuum—from core grasslands requiring preservation to transitional landscapes where alternative approaches may be more appropriate.

## Data Availability

The data can be visualized and accessed in **Google Earth Engine** via the following application: (https://wlfw-um.projects.earthengine.app/view/cropland-extent-map). The dataset is also available for download from **Zenodo**: https://zenodo.org/records/14607083. The codebase used in this work is openly available on **GitHub**: https://github.com/COYE-Coder/Annual-Cropland-Extent.

## CRediT author statement

**S. Carter**: Methodology, Software, Validation, Formal Analysis, Investigation, Data Curation, Writing - Original Draft, Writing - Review & Editing, Visualization; **S. Morford**: Conceptualization, Methodology, Software, Resources, Investigation, Writing - Original Draft, Writing - Review & Editing, Visualization, Supervision, Project administration; **J. Tack**: Conceptualization, Investigation, Writing - Review & Editing, Funding acquisition; **T. Lark**: Validation, Investigation, Writing - Review & Editing. **N. Uludere Aragon**: Validation, Investigation, Writing - Review & Editing; **B. Allred**: Methodology, Writing - Review & Editing; **D. Twidwell**: Writing - Review & Editing; **D. Naugle**: Conceptualization, Writing - Review & Editing, Supervision, Funding acquisition.

## Acknowledgements

The U.S. Fish and Wildlife Service supported this project through a grant administered by the Intermountain West Joint Venture and by the USDA Natural Resources Conservation Service (NRCS). We thank Kris Mueller for developing the grasslands chips and for assistance with figure production. The findings and conclusions in this paper are those of the authors and do not necessarily represent the views of the USDA NRCS or the U.S. Fish and Wildlife Service. The funding agency, which had no influence over the content’s review or approval provided funding for open access publication.

## Declaration of Generative AI Use

Google AI Studio (Model 1206 Experimental) was used to refine select sentences for clarity and conciseness. No generative AI was employed to create original content, prepare/edit figures, or analyze data. After using AI Studio, the author(s) reviewed and edited the material for accuracy and take(s) full responsibility for the published content.

## Supplemental Materials

### Supplemental 3.1 Model Training Process

1. *Pre-training (GLAD data only)*: We first pre-trained the model using only the synthetically generated GLAD labels for 25 epochs, where each epoch represents one complete pass through the entire training dataset. This allowed the model to learn general features of croplands across a broad spatial extent. We used a maximum learning rate of 0.0001.
2. *Encoder Freezing (GLAD data):* We then froze the weights of the encoder layers and trained the decoder layers for an additional 50 epochs using the same GLAD data. This approach, a form of transfer learning, leverages the pre-trained encoder’s ability to extract relevant features while allowing the decoder to specialize in cropland identification within our specific study area and prevent overfitting. We used a maximum learning rate of 0.001 with decay for this step.
3. *Fine-tuning (Chip data):* Next, we incorporated the manually annotated “chip” dataset to fine-tune the model’s performance in complex landscapes and address regional biases. Keeping the encoder weights frozen, we trained the decoder for 50 epochs with a maximum learning rate of 0.001 with decay. The high-quality chip data allowed the model to learn subtle variations in cropland characteristics that might not be captured in the synthetic GLAD labels.
4. *Full Training (Chip data)*: Finally, we unfroze all model weights and trained the entire network for 100 epochs using the high-quality chip dataset alone. This final training stage allowed the model to optimize all parameters and achieve favorable performance across disparate spatiotemporal domains. We increased the maximum learning rate to 0.005 with decay to facilitate further refinement of the model’s weights.

We used the Adam optimizer, known for its efficient performance in deep learning applications, and the Focal Tversky loss function, which addresses class imbalance in segmentation tasks and is particularly robust to false positives (Abraham and Khan, 2019). All training was performed using TensorFlow version 2.8 on an NVIDIA A100 Tensor Core GPU.

### Supplemental 3.2 Random Forest forested area mask

To address model errors of commission within the continental US study area, we developed a primary forest mask using a Random Forest (RF) model. The RF model was trained on a maximum composite of Normalized Difference Vegetation Index (NDVI), Gross Primary Productivity, winter and summer precipitation, Rangeland Analysis Platform (RAP) data, and National Land Cover Database (NLCD) labels. We created this custom model to improve forest and woodland detection in Native American reservations, where NLCD often underperformed. The model was trained on 500 randomly distributed NLCD-labeled points (forest and non-forest) and 50 additional points from areas where NLCD omitted primary forests (in total, 550 points). Using Google Earth Engine’s ***ee.Classifier.smileRandomForest()*** with 50 estimators, we achieved 93% accuracy on an unseen validation dataset. The resulting mask was applied to our dataset across the conterminous United States.

### Supplemental 3.3 Stratified sampling for reference points

Following the *Sample Analysis* methodology undertaken in Potapov et al. (2022), we defined 5 validation strata to evaluate model performance across space and time, providing a conservative estimate of model accuracy. These strata are: 1) *Stable croplands*; areas where cropland was predicted in at least 20 out of 26 images. 2) *Cropland gain*; regions predicted as cropland in 202 1 but not in 1996. 3) *Cropland loss*; Regions predicted as cropland in 1996 but not in 2021 (inverse of *cropland gain*). 4) *Possible errors*: areas where 5 or more images predicted cropland values between 0.2 5 and 0.75. 5) *Likely non-cropland*: areas where cropland was not predicted in at least 20 out of 26 images. Each stratum was sampled 54 times per year for a total of 7,020 reference points.

